# CRISPR-On-Beads: A Simple and Sensitive Approach for Bacterial DNA Detection

**DOI:** 10.1101/2025.05.02.651981

**Authors:** Jiaxiang Ye, Li Liu, Ruonan Peng, Fengjun Xu, Yujie Men, Ke Du

**Affiliations:** Department of Chemical and Environmental Engineering, University of California, Riverside, CA, 92521, USA; Department of Medical Oncology, Guangxi Medical University Cancer Hospital, Nanning 530021, China

## Abstract

CRISPR/Cas12a, recognized for its high efficiency and specificity in nucleic acid recognition, has found extensive applications in infectious disease sensing. Nonetheless, a significant challenge persists in devising a simple method for the rapid, convenient, and accurate visualization of detection results. In this study, we demonstrate an innovative and simple CRISPR-on-beads assay for visual detection of methicillin-resistant *Staphylococcus aureus* (MRSA). This approach combines isothermal target amplification, CRISPR/LbCas12a-mediated cleavage, and magnetic bead-based probe binding for signal amplification, enabling rapid and convenient identification of bacterial DNA. Our results show that this detection method exhibits both high sensitivity and specificity, with a detection limit as low as 30 CFU/μL. This newly developed biosensing approach can easily be integrated with a portable fluorescence microscope for medical diagnostics in resource-limited settings.

## 1. Introduction

Methicillin-resistant *Staphylococcus aureus* (MRSA) is a formidable pathogen capable of causing a wide spectrum of infections, ranging from superficial skin infections to life-threatening conditions such as sepsis and pneumonia[^1, 2^]. MRSA strains have been isolated from many foods and can spread throughout the food production chain, posing a serious threat to food safety. Moreover, the emergence and proliferation of MRSA, particularly in healthcare environments, present a significant challenge due to its resistance to β-lactam antibiotics, including methicillin^[1]^. Infections caused by MRSA are associated with significantly higher mortality rates compared to those caused by methicillin-susceptible strains, especially among immunocompromised individuals and those with underlying health conditions[^2, 3^]. Therefore, timely diagnosis and the implementation of effective treatment strategies are essential for reducing MRSA-related mortality.

Previous research has developed nucleic acid detection approaches for MRSA, utilizing quantitative real-time PCR (qPCR)[^4, 5^]. Although qPCR enables the rapid identification of resistance genes, it still requires specialized laboratories and advanced equipment, limiting its application in primary healthcare settings and remote areas. Recently, the Clustered Regularly Interspaced Short Palindromic Repeat (CRISPR)-associated nuclease (CRISPR/Cas) system has achieved remarkable progress in pathogen nucleic acid detection, particularly after the discovery of its collateral cleavage activity on adjacent single-stranded DNA (ssDNA)^[6]^. CRISPR/Cas-based biosensors have been effectively applied for the detection of bacterial and viral nucleic acids^[7-10]^. In addition, isothermal amplification methods such as loop-mediated isothermal amplification (LAMP)^[11]^ and recombinase polymerase amplification (RPA)^[12]^ have been incorporated into the CRISPR/Cas system to improve detection sensitivity[^13, 14^].

Fluorescence resonance energy transfer (FRET) DNA probes have been widely used to enable the collateral cleavage activity of CRISPR systems[^15, 16^]. Even though highly sensitive and simple, they present challenges such as high background and quenching issues. To address these problems, nanomaterials with higher and more stable fluorescence signals, such as quantum dots have been incorporated into CRISPR assays. But it increases the complexity of the assay^[17]^. Therefore, a practical approach is to concentrate the cleaved FRET probes on an extended solid surface to increase the fluorescence intensity^[18]^; thus, the signal can easily be resolved by various imaging techniques.

In this study, we introduce a straightforward and sensitive CRISPR-on-Beads assay for the rapid detection of the MRSA resistance gene, *mecA*, with a detection limit of 30 CFU/µL. A modified single-stranded DNA (ssDNA) reporter tagged with biotin, Cy5, and a quencher was designed for CRISPR trans-cleavage. Following RPA and the CRISPR reaction, the cleaved ssDNA is captured by streptavidin-coated beads and can easily be quantified using a fluorescence microscope. Without bead-based collection and concentration, the cleaved probes remain undetectable under the microscope, highlighting the importance of solid-surface concentration. Moreover, each test requires only ∼3 µL of sample, making it well-suited for handling low-volume patient specimens. This simple and sensitive bead-based assay opens new possibilities for point-of-care detection of various infectious diseases.

## 2. Experimental Methods

### 2.1. Reagents

The matched target sequence was selected from the *mecA* gene in the MRSA strain. The target sequence, RPA primers, CRISPR-RNA (crRNA), biotin/Cy5-quencher (BC-Q) single-stranded DNA (ssDNA) probes, AsCas12a, and LbCas12a were obtained from Integrated DNA Technologies (IDT), with detailed specifications of the synthetic oligonucleotides provided in **Table 1**. Lysozyme, PureLink™ Genomic DNA Mini Kits (Catalog No. K1820-00, Invitrogen™), and Dynabeads MyOne Strepavidin C1 magnetic beads were sourced from ThermoFisher Scientific. The TwistAmp® Basic Kit was acquired from TwistDx™. Additionally, NEBuffer™ r2.1 buffer was procured from New England Biolabs.

**Table 1.**
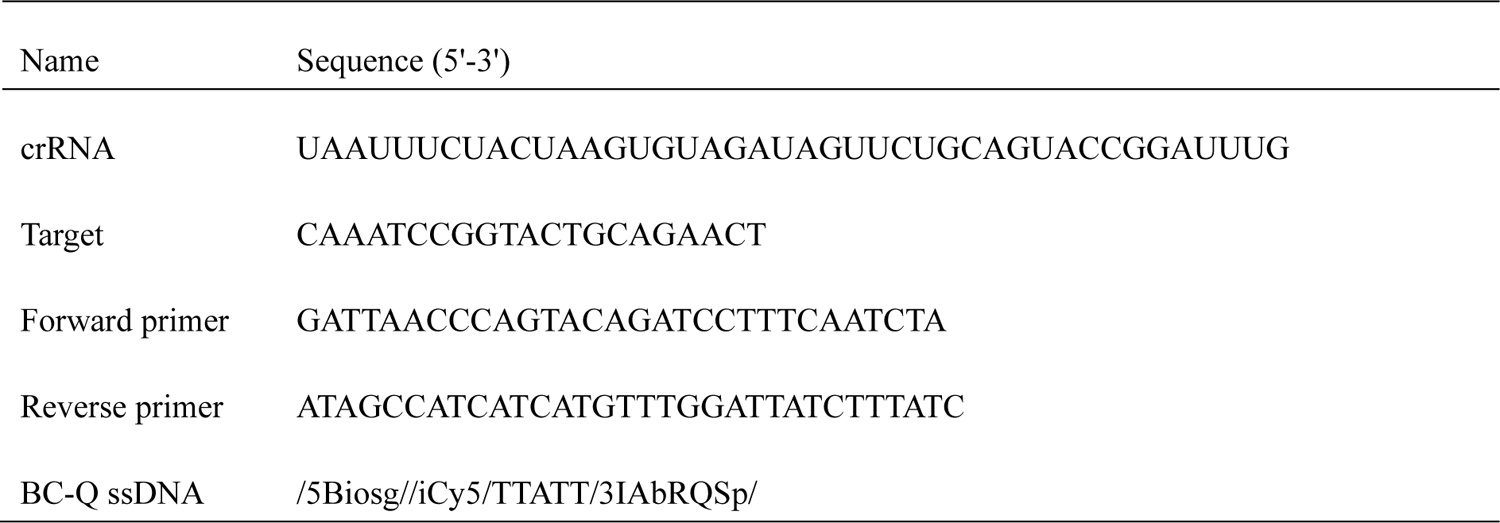
List of synthetic oligos sequence used in this study.

### 2.2. Bacterial culture

Methicillin-resistant *Staphylococcus aureus* (MRSA), wild-type *Staphylococcus aureus* (SA), and *E. coli* strains MG1655, DH5α, and TOP10 were obtained from laboratory stock. The bacterial strains were cultivated overnight in tryptic soy broth (MilliporeSigma) at 37°C with agitation at 200 rpm in a bacterial incubator. Subsequently, 1 mL of the bacterial suspension was centrifuged at 8,000 × g for 5 minutes to pellet the cells. The pellet was then washed twice by resuspending in 1 mL of phosphate-buffered saline (PBS) to thoroughly remove any residual culture medium. The bacteria concentration was determined by the colony-forming unit (CFU) counting method using standard tryptic soy agar (TSA) plates. For subsequent experiments, the harvested bacteria were serially diluted in PBS.

### 2.3. Nucleic Acid Extraction

Nucleic acid extraction was performed through bacterial lysis followed by DNA purification using PureLink™ Genomic DNA Mini Kits. Following the manufacturer’s instructions for the Genomic DNA Mini Kits, gram-positive bacteria (*MRSA* and *SA*) were initially incubated in 180 μL of lysozyme digestion buffer (25 mM Tris-HCl, pH 8.0, 2.5 mM EDTA, 1% Triton X-100) containing 20 mg/mL lysozyme, while gram-negative bacteria (*E. coli*) were incubated solely in the genomic digestion buffer. Subsequently, 20 μL of proteinase K was added to gram-positive cells, and 20 μL of RNase A was added to gram-negative cells. For DNA purification, both gram-positive and gram-negative cells followed an identical spin column-based centrifugation procedure. Finally, the purified total DNA was eluted in 30 μL of elution buffer.

### 2.4. RPA Amplification

RPA reaction was performed using the TwistAmp® Basic Kit following the manufacturer’s instructions. A mixture containing 2.4 μL each of forward and reverse primers (10 μM), 29.5 μL of rehydration buffer, and 11.2 μL of nuclease-free water was added to the enzyme pellet. Subsequently, 2 μL of extracted total DNA and 2.5 μL of MgOAc (280 mM) were added to reach a total reaction volume of 50 μL, followed by incubation at 39°C for 20 minutes.

### 2.5. CRISPR-Cas12a-Based Cleavage

Initially, 62.5 nM crRNA was complexed with 50 nM AsCas12a or LbCas12a at room temperature for 10 minutes. Subsequently, the optimal BC-Q ssDNA probes, 1× binding buffer, 13.25 μL nuclease-free water (IDT, Inc.), and 2 μL of DNA sample were combined with the As or LbCas12a-crRNA complex, yielding a final reaction volume of 20 μL. All DNA samples used in the assay consisted of either synthetic target sequences from IDT or RPA-amplified products of total DNA extracted from MRSA. The reaction mixture was incubated at 37°C for 30 minutes. After incubation, Cy5 fluorescence signals (excitation wavelength: 645 nm, emission wavelength: 665 nm) were quantified from the mixture prior to Cy5-bound magnetic beads and characterized using an imaging reader (Agilent BioTek Cytation 5) to confirm fluorescence microscopy detection.

### 2.6. Fluorescence Microscopy Imaging

The supernatant of the 10 μL magnetic bead solution (Dynabeads MyOne Strepavidin C1) was discarded by isolating the streptavidin-coated magnetic beads with a magnetic rack (NEBNext magnetic separation rack, New England BioLabs, Inc.). The magnetic beads were washed three times with PBS and resuspended in 10 μL of PBS. The washed magnetic beads were then transferred to a new 0.6 mL PCR tube, and 10 μL of the cleaved CRISPR products were added to the magnetic beads at room temperature for 30 minutes. After incubation in a slow shaker, the magnetic bead-Cy5 conjugates were washed and resuspended in 20 μL of PBS for later use. Three microliters from each tube were placed onto a microscope slide, and fluorescent images were captured using a Leica DMi8 inverted microscope. The mean fluorescence intensity of these images was quantified with Leica LAS X software.

### 2.7. Statistical analysis

Data analysis was performed using GraphPad Prism 8.1 (GraphPad, San Diego, CA, USA) based on at least three independent experiments. Continuous data were represented as mean ± standard deviation (mean ± SD). Statistical comparisons for normally distributed data were conducted using a two-tailed t-test, unless otherwise specified. A *p*-value of less than 0.05 was considered statistically significant.

## Results

The detection mechanism of MRSA via the CRISPR-on-beads system is depicted in **Figure 1**. The Cas12a/crRNA complex specifically recognizes and binds to the target DNA, activating the trans-cleavage function of Cas12a on BC-Q ssDNA. In this study, the BC-Q ssDNA is labeled with a fluorescent dye (Cy5) linked to biotins and a quencher group (Iowa Black® RQ-Sp). BC-Q ssDNA serves as both a fluorescent reporter and a molecule that binds to streptavidin-coated magnetic beads, forming magnetic bead-Cy5 conjugates. Upon the introduction of magnetic beads into the cleaved reaction products, the fluorescence of the magnetic bead-Cy5 conjugates became detectable under a fluorescence microscope. This approach facilitates both qualitative and quantitative detection of MRSA through the analysis of magnetic bead-Cy5 conjugates. The cleavage kinetics of CRISPR/Cas12a were systematically analyzed using our BC-Q probe to optimize Cas12a protein variants for downstream applications. The corresponding Michaelis-Menten fitting curves[^6, 19^] for cleavage activity were generated using the mean enzyme velocity (V) and substrate concentration (BC-Q ssDNA or target DNA) with GraphPad Prism 8.1. We first assessed the trans-cleavage activity of LbCas12a and AsCas12a over time across target concentrations ranging from 0.05 to 3 ng/µL, with the BC-Q probe fixed at 2,500 nM. As shown in **Figure 2a**, LbCas12a achieved fluorescence intensities of ∼10,000 counts at target concentrations above 0.75 ng/µL, whereas AsCas12a reached only ∼2,000 counts, indicating superior trans-cleavage efficiency of LbCas12a (**Figure 2b**). Further kinetic analysis at varying probe concentrations (**Figure 2c** and **2d**) showed that LbCas12a reached ∼25,000 counts at 5,000 nM after 30 minutes, while AsCas12a remained below 3,000 counts under the same conditions. The plot of mean reaction rate (V) versus target concentration (**Figure 2e**) exhibits strong linearity for both enzymes, confirming the higher catalytic efficiency of LbCas12a. Based on these results, LbCas12a was selected for subsequent CRISPR-on-beads experiments.

**Figure 1.**
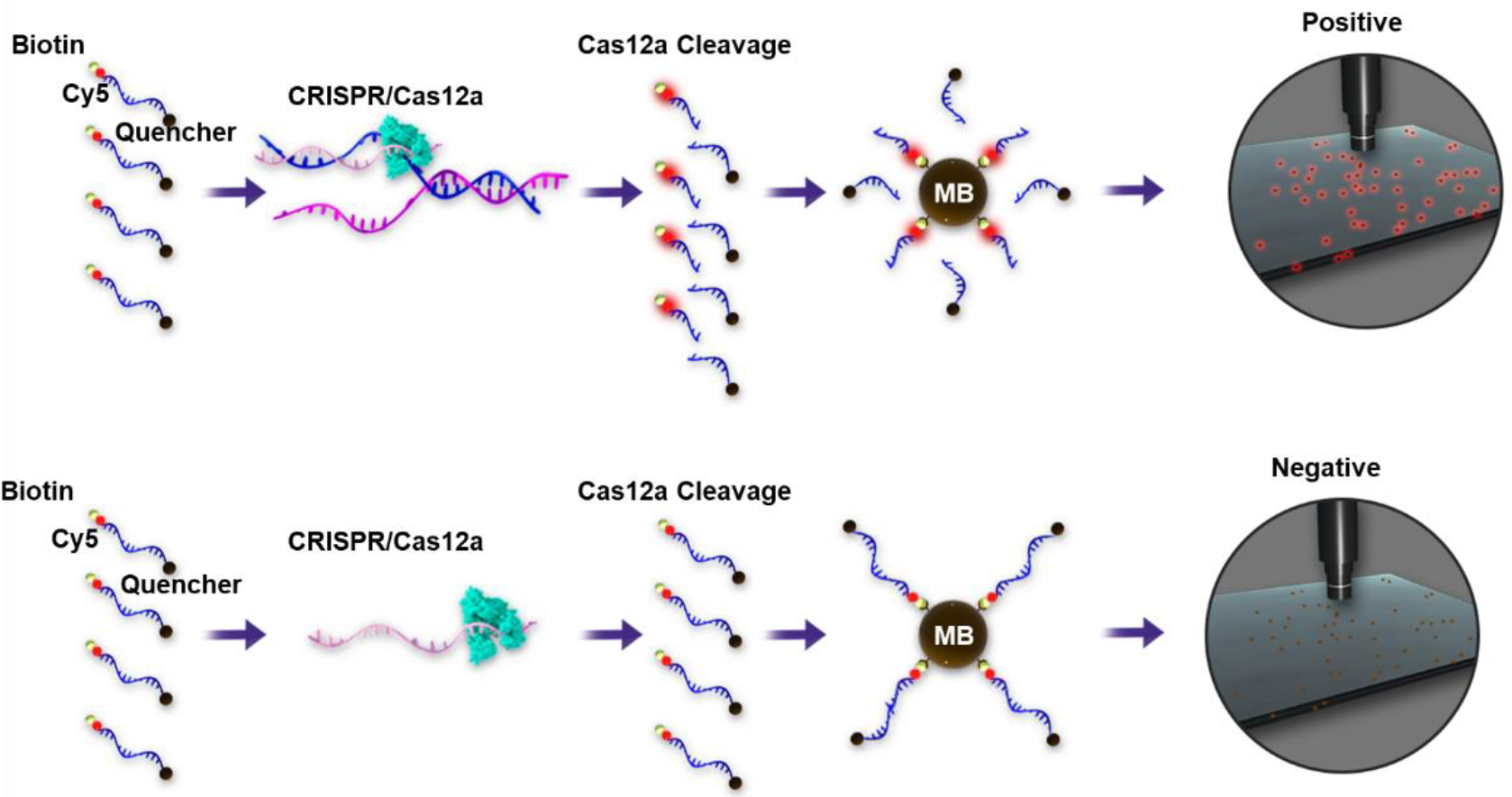
Schematic illustration of the bacteria detection via CRISPR-on-beads. The activated CRISPR complex cleaves the BC-Q probes and then be immobilized on the magnetic beads for imaging. Without target activation, the fluorescence dye is quenched on the beads without showing fluorescence signals.

**Figure 2.**
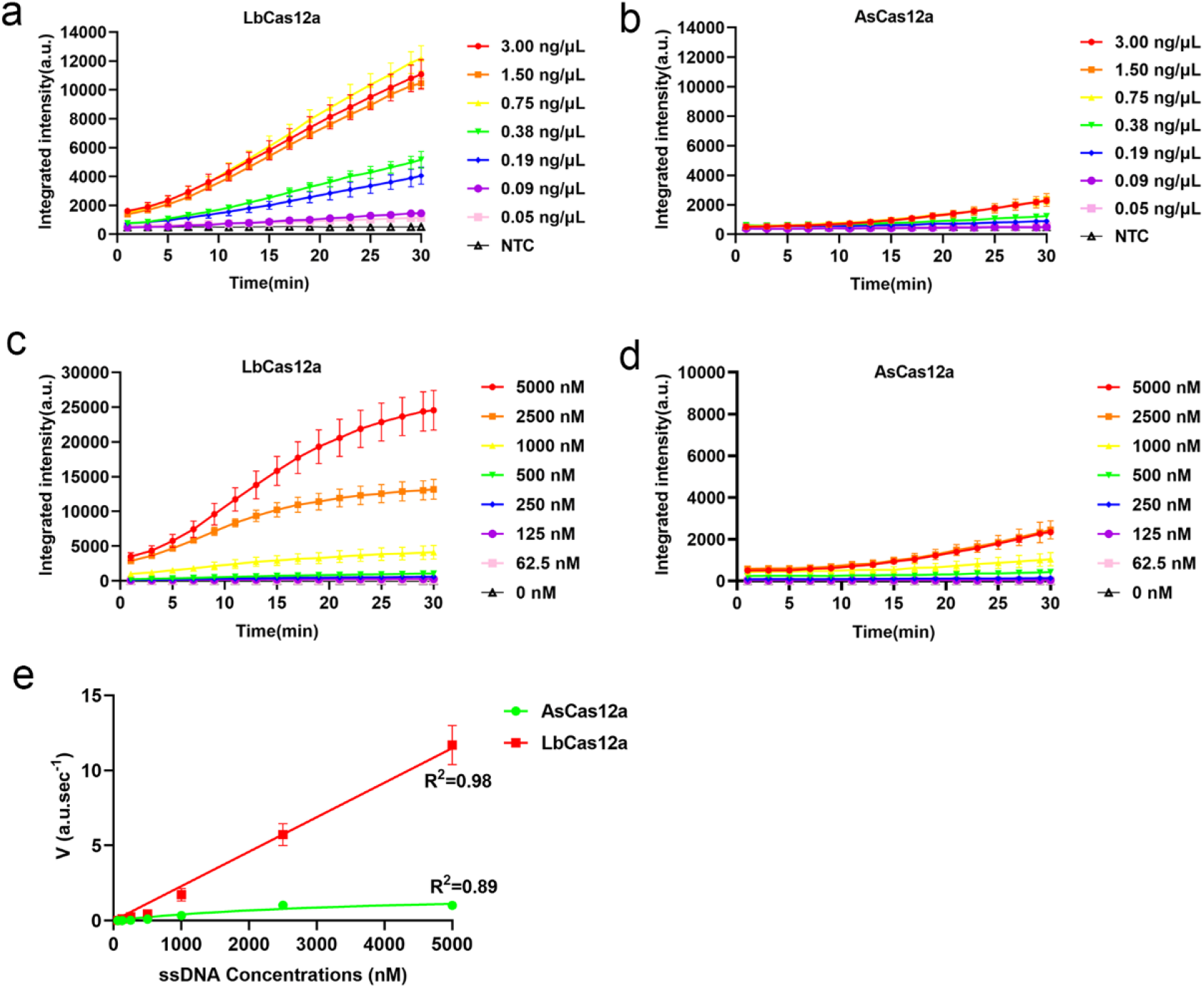
Interval plots showing trans-cleavage velocity over time for (a) LbCas12a and (b) AsCas12a at varying MRSA concentrations (0.05 to 3.00 ng/µL), measured over 30 minutes. Interval plots for (c) LbCas12a and (d) AsCas12a at varying ssDNA probe concentrations (0 to 5,000 nM). (e) Michaelis-Menten curves illustrating the trans-cleavage kinetics of AsCas12a and LbCas12a.

To establish a CRISPR-on-beads protocol, varying concentrations of BC-Q ssDNA (0–5 µM) were incubated with 1.00 ng/μL of synthetic target DNA at 37°C for 30 minutes. The fluorescence intensity of the cleaved products was first measured using an image reader prior to bead binding (**Figure 3a**). Magnetic beads were then introduced, and the samples were imaged using a microscope. While 5 µM BC-Q ssDNA yielded the highest fluorescence intensity before bead binding, the clearest fluorescence signal post-binding was observed at 2.5 µM (**Figure 3b**). As a result, 2.5 µM BC-Q ssDNA was selected for subsequent experiments. Next, magnetic bead concentrations of 0, 2.5, 5, and 10 μg/μL were evaluated using cleaved CRISPR products generated with 2,500 nM BC-Q ssDNA and 1.00 ng/μL synthetic target DNA to optimize bead–Cy5 conjugation. As shown in **Figure 3c** and **3d**, 5 μg/μL of magnetic beads produced the strongest fluorescence signal and was therefore selected for future experiments.

**Figure 3.**
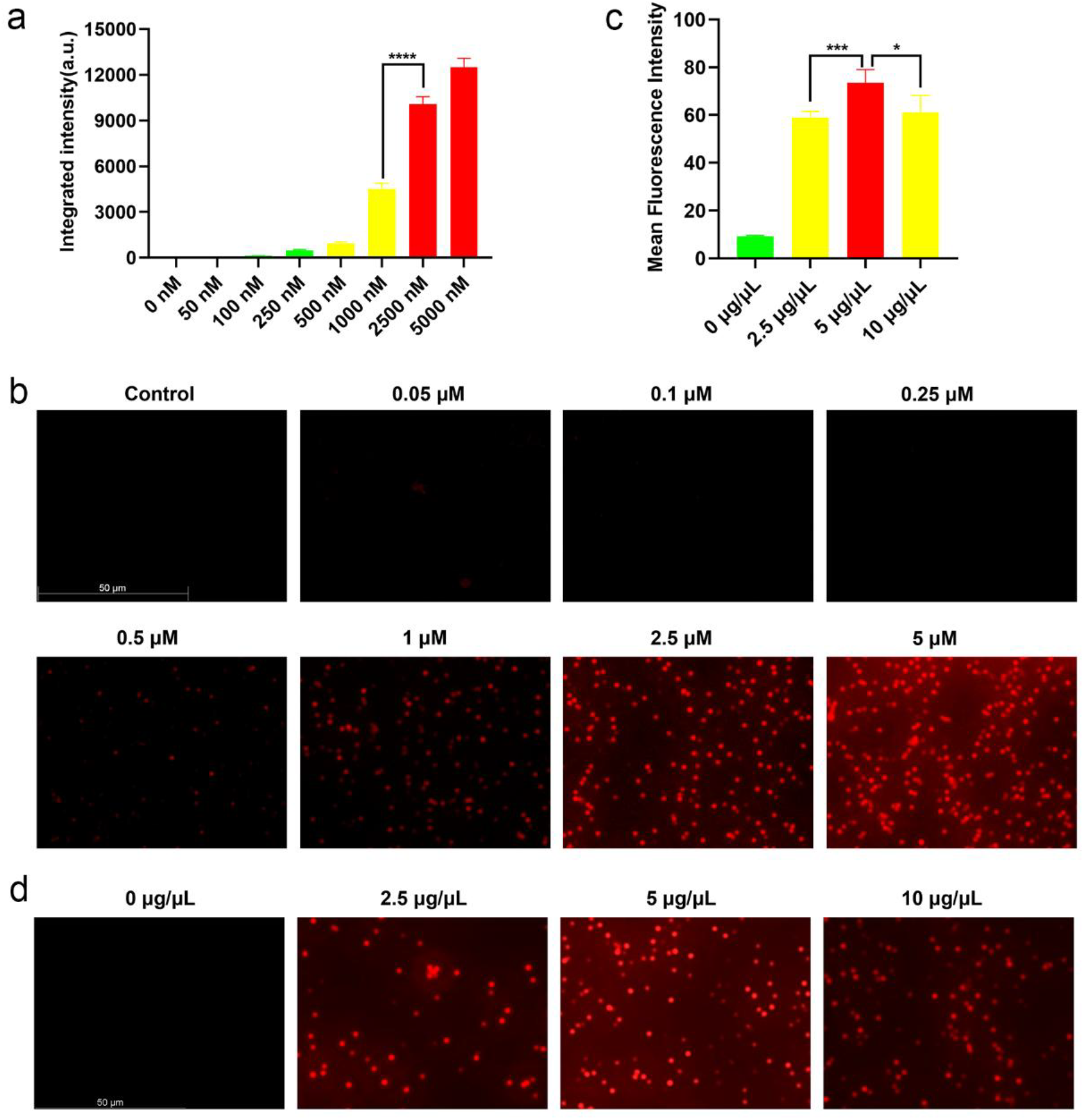
(a) Quantified fluorescence signals from CRISPR/LbCas12a reactions at different BC-Q ssDNA concentrations measured by an imaging reader before Cy5 binding to magnetic beads (Ex/Em = 645/665 nm). (b) Fluorescence microscope images of magnetic bead-Cy5 conjugates at varying concentrations of BC-Q ssDNA in CRISPR/LbCas12a reactions. Red dots indicate Cy5 conjugated beads. (Scale bar: 50 μm). (c) Mean fluorescence intensity of magnetic bead-Cy5 conjugates at different concentrations of magnetic beads (Ex/Em = 645/665 nm). (d) Fluorescence microscope images of magnetic bead-Cy5 conjugates at different concentrations of magnetic beads with fixed volume (10 μL) of cleaved reaction products. Data are presented as mean ± SD (\**p* < 0.05, \*\*\**p* < 0.001, \*\*\*\**p* < 0.0001, Student’s t-test).

The performance of the CRISPR/Cas12a-based detection strategy for synthetic target DNA is shown in **Figure 4**. Fluorescence microscope images of magnetic bead-Cy5 conjugates at varying concentrations of synthetic target DNA (0.05 to 0.75 ng/µL) are displayed in **Figure 4a**, while the corresponding mean fluorescence intensity is shown in **Figure 4b**. To confirm the fluorescence microscopy detection, the quantified fluorescence signals from CRISPR/LbCas12a reactions prior to Cy5 binding to the magnetic beads were measured using an imaging reader, as shown in **Figure 4c**. Collectively, these results indicate that we can detect synthetic target DNA with a concentration of 0.09 ng/μL with our CRISPR-on-beads assay. To enhance the detection sensitivity while maintaining operational simplicity, RPA was employed for target amplification, while the CRISPR/Cas12a system was utilized for signal readout. Compared to direct detection without RPA, the detection limit in the RPA+CRISPR/LbCas12a reactions improved by at least four orders of magnitude in MRSA concentration. As shown in **Figure 5**, the RPA-CRISPR/LbCas12a approach achieves a detection limit of 3×10^4^ CFU/mL for MRSA.

**Figure 4.**
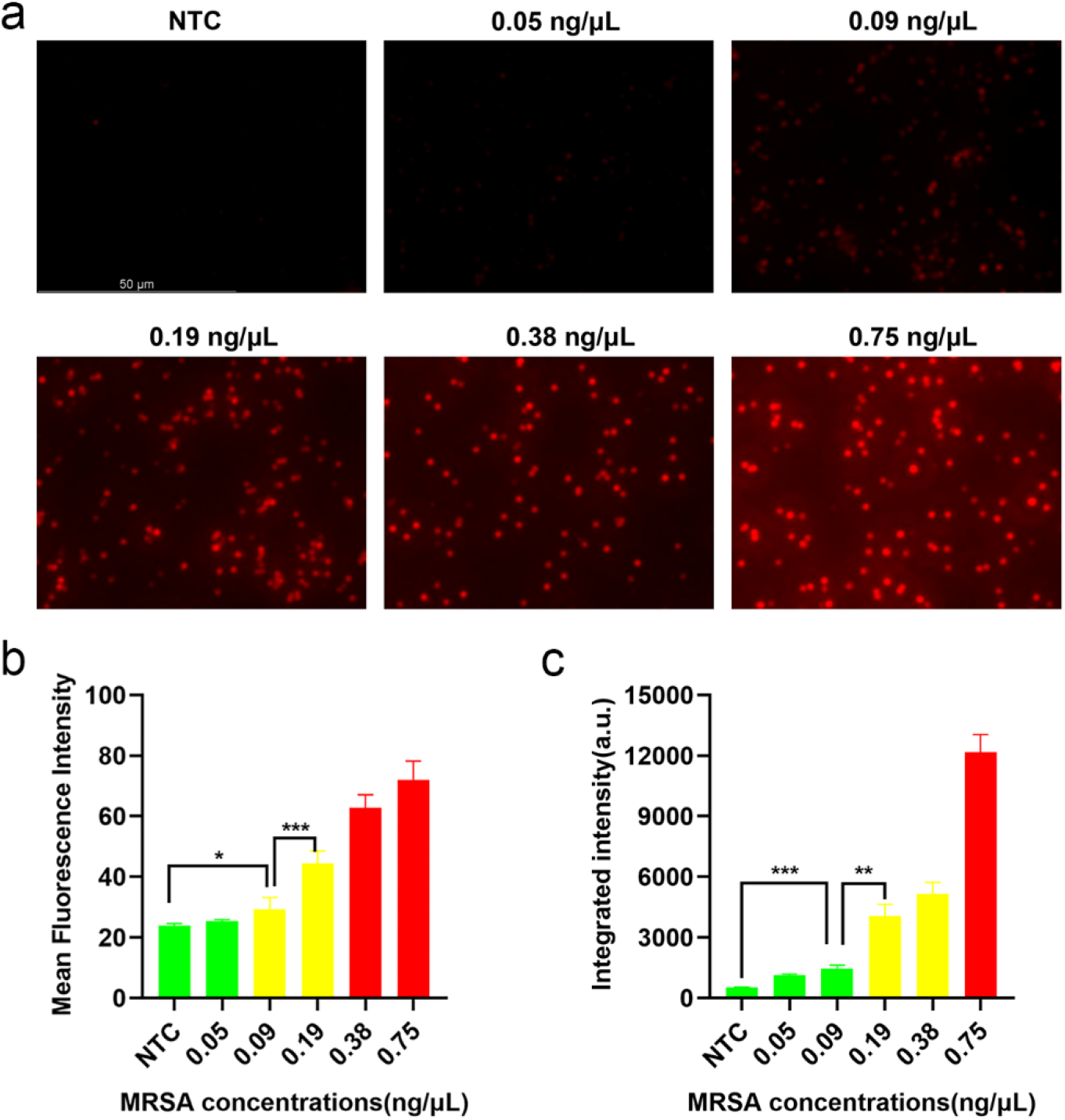
(a) Fluorescence microscope images of magnetic bead-Cy5 conjugates at varying concentrations of synthetic target DNA. Red dots indicate Cy5 labeled beads (Scale bar: 50 μm). (b) Mean fluorescence intensity of the images in (a). (c) Quantified fluorescence signals from CRISPR/LbCas12a reactions at different synthetic target DNA concentrations, measured by an imaging reader before Cy5 binding to magnetic beads (Ex/Em = 645/665 nm). Data are presented as mean ± SD (\**p* < 0.05, \*\**p* < 0.005, \*\*\**p* < 0.001, Student’s t-test). NTC represents the no-template control.

**Figure 5.**
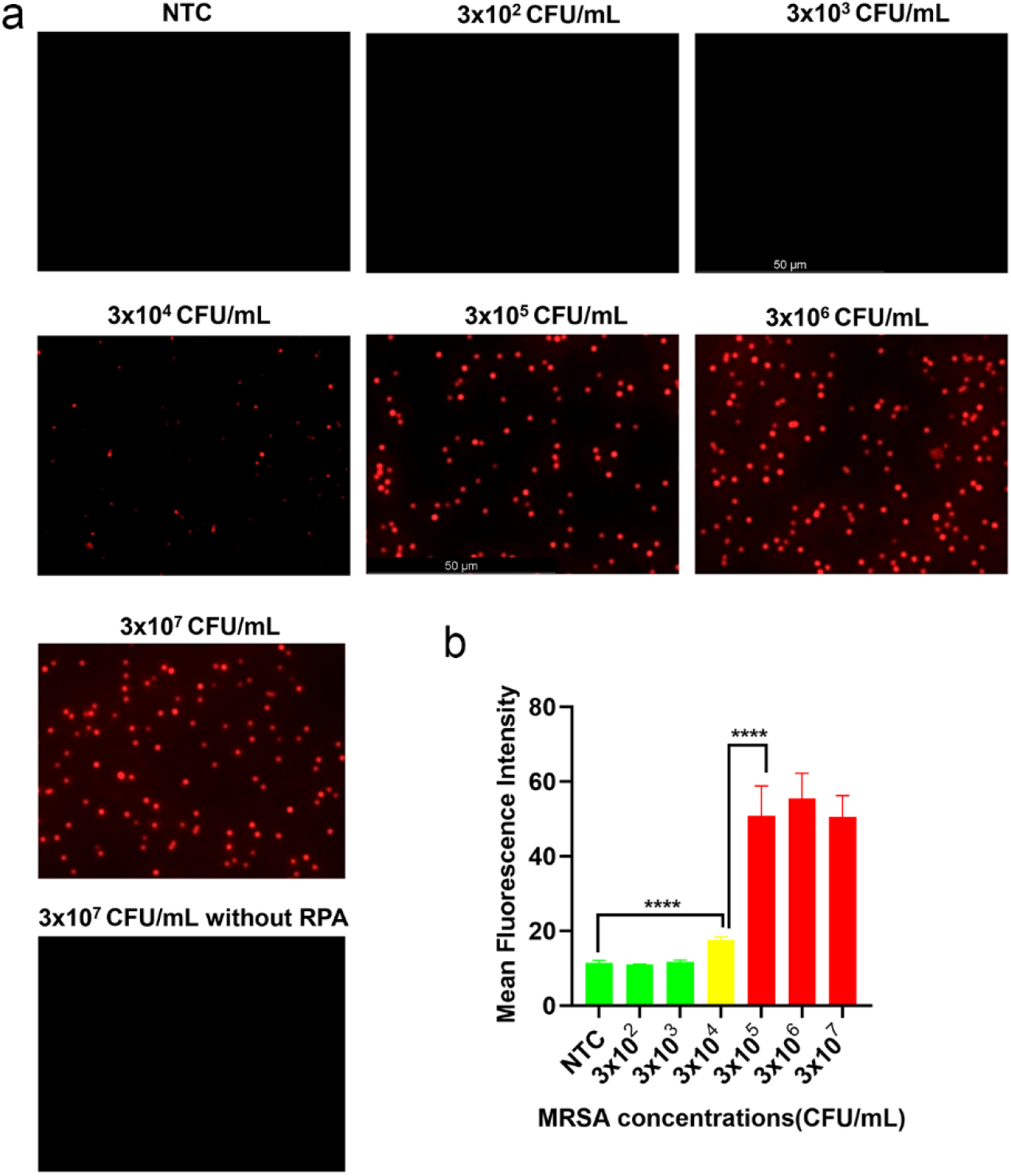
(a) Detection limit of CRISPR-on-beads determined by fluorescence microscope images. The MRSA target concentration ranging from 300 to 3×10^6^ CFU/mL. The Red dots indicate Cy5 labeled beads (Scale bar = 50 μm). (b) Mean fluorescence intensity of the images in (a). Data are presented as mean ± SD (\*\*\*\**p* < 0.0001, Student’s t-test). NTC represents the no-template control.

Two μL of total DNA, extracted from various bacterial sources at a uniform concentration of 3×10^7^ CFU/mL, was utilized for RPA. The specificity of the RPA-CRISPR/Cas12a detection strategy for nucleic acids was then assessed by introducing 2 μL of RPA-amplified DNA into 18 μL of CRISPR/Cas12a detection reagents. DNA extracted from five lysed bacterial strains, including *MRSA*, wild-type *SA*, and *E. coli* strains *MG1655, DH5α*, and *TOP10*, was added to the assay. As illustrated in **Figure 6**, the LbCas12a-based detection system exhibits robust specificity for MRSA, with no detectable signal in other bacterial samples.

**Figure 6.**
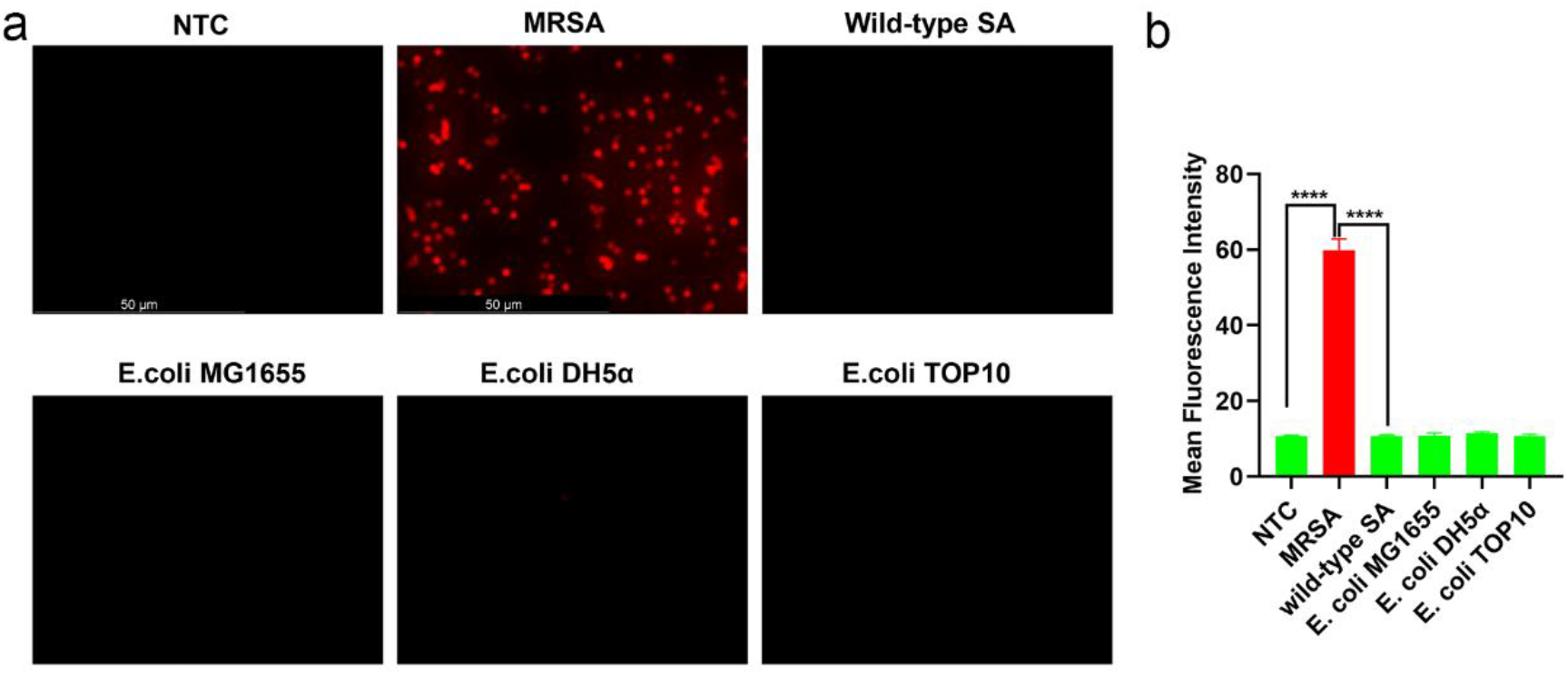
(a) Specificity test of the CRISPR-on-beads strategy. The fluorescence microscope images of magnetic bead-Cy5 conjugates for various bacteria targets with only MRSA showing positive signals (Scale bar: 50 μm). (b) Mean fluorescence intensity of the images in (a). Data are presented as mean ± SD (\*\*\*\**p* < 0.0001, Student’s t-test). NTC represents the no-template control.

## Discussion

In this study, we integrate sequence-specific RPA with CRISPR-Cas12a to detect the MRSA resistance gene *mecA*, enabling precise differentiation between antibiotic-resistant and wild-type strains. Both the RPA and CRISPR reactions are carried out under isothermal conditions at 37°C, eliminating the need for complex equipment such as thermal cyclers. Additionally, we compared the kinetic performance of two Cas12a variants, LbCas12a and AsCas12a, and found that LbCas12a demonstrates superior activity, resulting in enhanced overall detection sensitivity.

Additionally, the concentration of the BC-Q probe in the detection system was optimized, with results indicating that 2,500 nM is the optimal concentration. Although higher probe concentrations initially produced stronger fluorescence signals prior to bead conjugation, the resulting immunofluorescence images lacked clarity. This may be due to excessive reaction product formation at elevated probe concentrations, leading to saturation of the magnetic beads and fluorescence overexposure. Therefore, 2,500 nM was selected as the optimal ssDNA probe concentration for subsequent assays, balancing image clarity under fluorescence microscopy with cost-effectiveness in detection.

We developed a more practical and effective alternative to focusing FRET probes on solid surfaces, magnetic beads to amplify fluorescence intensity. This method allows the fluorescent signal to be more easily resolved using imaging techniques. By utilizing magnetic beads, the fluorescently labeled probes throughout the solution volume are concentrated into a small detection area, which greatly increases the sensitivity of the assay^20, 21^]. This concentrated fluorescence increases the signal-to-noise ratio and improves the reliability and detection limit of the assay^[22]^.

In addition, the use of magnetic beads offers the dual advantage of not only enhancing the fluorescence intensity but also facilitating the separation of the signal from the background noise. Magnetic beads, in combination with fluorescently labeled probes, utilize magnets to selectively attract and separate labeled particles while non-specific background components remain in the supernatant. The supernatant is then removed, leaving a purified and concentrated sample for analysis. This streamlined process simplifies workflow, increases assay sensitivity, and minimizes background interference, making it an attractive solution for enhancing FRET-based CRISPR assays. The results indicate that the MRSA visual detection method displays both high sensitivity and specificity, achieving a detection limit as low as 30 CFU/μL. Building on current detection outcomes, integrating this method with a portable fluorescence microscope will significantly enhance its applicability in field settings and resource-limited environments, including rural and remote regions[^23, 24^].

The developed method could be further optimized in the following aspects to target complex sample matrices and various antibiotic resistance mechanisms. Although cultured MRSA strains were successfully detected, the application to real samples, such as food matrices and patient blood, may also require the development of an efficient and user-friendly DNA extraction procedure. Additionally, MRSA resistance mechanisms are complex and involve multiple genes, including *mecA, mecC, blaZ*, and *vanA*^[1, 25, 26]^. Therefore, the development of a multiplexed CRISPR-on-beads assay would be highly beneficial for simultaneously identifying these resistance genes, ultimately contributing to improved monitoring of antibiotic resistance in both food safety and healthcare settings.

## Acknowledgements

This project is supported by SoCal OASIS™ Internal Funding Awards.

